# Pseudouridylation of rRNA by specific snoRNA disrupts ribosomal machinery and consequently affects metabolism, longevity and neurodegeneration

**DOI:** 10.64898/2026.04.17.719250

**Authors:** Théo Gauvrit, Pauline Minquilan, Virginie Marchand, Yuri Motorin, Jean-René Martin

## Abstract

In our society, ageing, longevity, and neurodegenerative diseases are major concerns of public health. Recently, in Drosophila, we have identified a new cluster of three snoRNAs, including *jouvence*, and showed that each of them affect longevity and neurodegeneration. As these snoRNAs are required in the epithelium of the gut, these results point-out a causal relationship between the epithelium of the gut and the neurodegenerative lesions through the metabolic parameters, indicating a gut-brain axis. Here, we demonstrate that each snoRNA pseudouridylates a specific site on ribosomal-RNA, which consequently affects the amount of ribosomes as well as the translational efficacy. Moreover, using TRAP experiment assay, we also show that these lacks of pseudouridylations modify the translation of specific genes involved in lipid metabolism. Consequently, these lead to a chronic deregulation of trigycerides and sterols levels, whose correlate to an increase of neurogenerative lesions in old flies, as well as to a modification of longevity.

## Introduction

The increasing life expectancy in our society is accompanied by a growing number of persons suffering from neurodegenerative and metabolic disorders, which represent a major problem of public health. These disorders results from complex biological processes of accumulation of damages at molecular, cellular, tissue, organ, and whole organism levels (Singh et al., 2019; Fontana et al., 2010). This high multi-levels complexity generally makes it difficult to precisely decipher the genetic and molecular origins of the cause, as to where and when they occurred. Up to now, these mutiple ageing causes have been resumed in 12 hallmarks (López-Otín et al., 2013; 2023). Thus, several genes and signaling pathways involved in the determination of ageing, neurodegeneration, and lifespan have been identified (López-Otín et al., 2016; 2023; Gems & Partridge, 2013; Broughton et al., 2005; Owusu-Ansah & Perrimon, 2014). Although the insulin signaling pathway has been the most investigated (Giannakou & Partridge; 2007; Broughton et al., 2005; Biglou et al., 2021), several other studies have also underlined the role of lipid metabolism (Johnson & Stolzing, 2019; Liu et al., 2015), such as the fatty-acid synthesis, triglycerides, and sterol (including the cholesterol), both in Drosophila (Jing & Behmer, 2020; Tschäpe et al., 2002), as well as in mammals (Saher, 2023). For example, the NPC1a causes cholesterol aggregation and age-progressive neurodegeneration (Phillips, et al., 2008), while the NPC1b gene promotes sterol absorption from the midgut epithelium (Voght et al., 2007). Moreover, NPC type C-2 genes family control sterol homeostasis and steroid biosynthesis, and has been used as a model of human neurodegenerative diseases (Huang et al., 2007). Finally, more recently, several microRNAs have also been shown to affect ageing and neurodegeneration (Eacker et al., 2009; Liu et al., 2012; Kato & Slack, 2013; Abe & Bonini, 2013; Soulé et al., 2020; Soulé & Martin, 2020; Al Issa et al., 2026). In that way, we have recently identified and characterized the role of a new cluster of three snoRNAs including jouvence, in the control of longevity, neurodegeneration and their relationship to lipids and sterol metabolism (Al Issa et al., 2026). Although we have shown that these three snoRNAs are principally required in the epithelium of the gut, several questions about their precise molecular mechanisms still remain to be elucidated.

Here, using different and complementary genetic, genomic, molecular, biochemical, cellular, and physiological approaches, in a context of whole organismal level, we first demontrate that each snoRNAs (jouvence, sno-2, and sno-3) pseudourydilates a specific site on ribosomal-RNAs, from which, two of them where not yet associated to a given snoRNA. To go further, we show that these defects in pseudouridylation perturb the ribosomes machinery. Indeed, by targetting a GFP-tagged ribosomal protein, we show that the amount of ribosomes is reduced. Moreover, using the puromycin incorporation, we show that the proteins synthesis rate is also reduced. In addition, using the TRAP (Translational Ribosomes Assay Purification), by targetting a Flag-tagged ribosomal protein, we demonstrated that the translation of some selected precific genes is affected, indicating that specific pseudouridylation of rRNA leads to a very selective defect in translation of some genes. To support these last results, we show by RT-qPCR that specific limiting genes involved in lipids and sterol metabolism are deregulated, which consequently leads to a deregulation of the overall triglycerides and sterol (cholesterol). Finally, to highlight the biological significance of these highly specific molecular defects, we show that the CRISPR/Cas9 mutation of each snoRNA affects fly longevity and neurodegeneration.

## Results

### Genomic Map of CRISPR/Cas9 mutants of each snoRNAs

We have recently characterized a new cluster of three small nucleolar RNA (snoRNA) including *jouvence*, in Drosophila, and showed that their deletion reduces lifespan, while their specific targetted rescue and/or overexpression within the epithelium of the gut is sufficient to increase and rescue the lifespan (Soulé et al., 2020; Al Issa et al., 2026). These demonstrations were mainly based on the simultaneous deletion (F4) of these three snoRNAs, in which we have rescued each of them individually or all together through the targetted P[Gal4] system. Although informative, this genetic approach does not inform us about their fine regulation neither on their molecular mechanisisms respectively. Here, in the aim to specifically disrupt a single snoRNA and let the two other one’s intact, we have generated a CRISPR/Cas9 mutant for each of them (Figure 1a). We use the replacement strategy of short sequence to disrupt the H-box, which is known to be essential for the fonctionaly of each snoRNA (Kiss, 2002). The replacement sequences are shown in Figure 1b.

**Figure 1.**
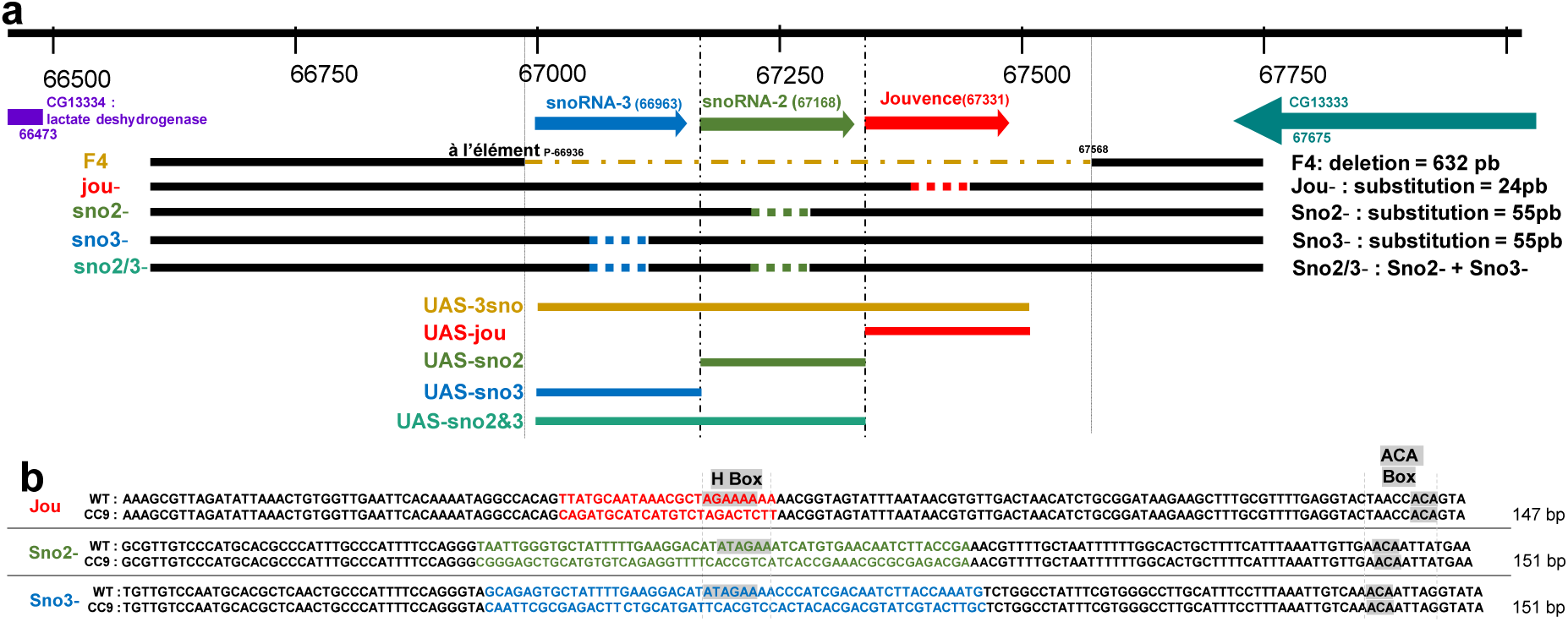
Molecular map of the P[Gal4] locus, and snoRNAs mutant lines. **a)** Genomic map of the 4C locus. The snoRNA:Ψ28S-1153 (jouvence), as well as the two other upstream snoRNAs (sno-2 = snoRNA:2R:9445205 and sno-3 = snoRNA:2R:9445410), are in the reverse orientation of the two encoding genes: CG13333 and CG13334. The deletion (F4) of 632 bp (caramel/brown dotted line) includes the three snoRNAs. Below are the four CRISPR-Cas9 mutant lines with the substitution (dotted line) in the H-box of the snoRNAs. Below are five of the UAS-line expressing part of the locus used in this study. **b)** Sequences of the three snoRNAs. For each snoRNA, the first line represents the wild type sequence, while the second represents the CRISPR/Cas9 (CC9) mutated sequence with the substitution in color. Highlighted in grey are the H-Box (when unmutated) and the ACA Box of each snoRNAs.

### The mutation of each snoRNA affects the expression level of the two others

To determine if the mutation caused by the replacement of each snoRNA locally affects the genetic locus and the expression level of the two others, we have performed RT-qPCR (Taqman) on each snoRNAs CRISPR/Cas9 mutants, both in young and old flies (Figure 2a,b). Intrigingly, although the replacement sequence was designed to have exactly the same number of nucleotides than the original sequence, the mutation in each snoRNA affects the expression level of the two others, indicating an interrelated and tight regulation of each of them at this locus (Figure 2c). Briefly, for *jouvence*, it seems that the sno-2 and the sno-3, which are both located upstream of *jouvence*, are essential for its expression, since jou is importantly reduced in sno-2 and sno-3 mutants, and almost absent in sno-2&3 mutant, suggesting that sno2 and sno3 could represent promoter-like sequence for *jouvence*. However, in contrast, the mutation of jou, located downstream to the sno-2 and sno-3, also affects their expression (an increase), suggesting a feedback control or a more complex regulation, as perhaps the processing of a longer pre-transcript, which remains to be determined. Even more surprisingly, the mutation of the sno-2, located between the sno-3 and jou, upregulates the expression of the sno-3, while similarly, the mutation of the sno-3, located upstream to the two others, also upregulates the sno-2, but decreases the expression level of jou. Finally, using the Gene-Switch system (Mex-GS) to conditionnaly rescue in adulthood each snoRNA in their own snoRNA mutant, do not rescue the expression level of the two others (Figure 2d,e,f), suggesting that there are no transcriptional retro-control, but rather a post-transcriptional regulation at the RNA level.

**Figure 2.**
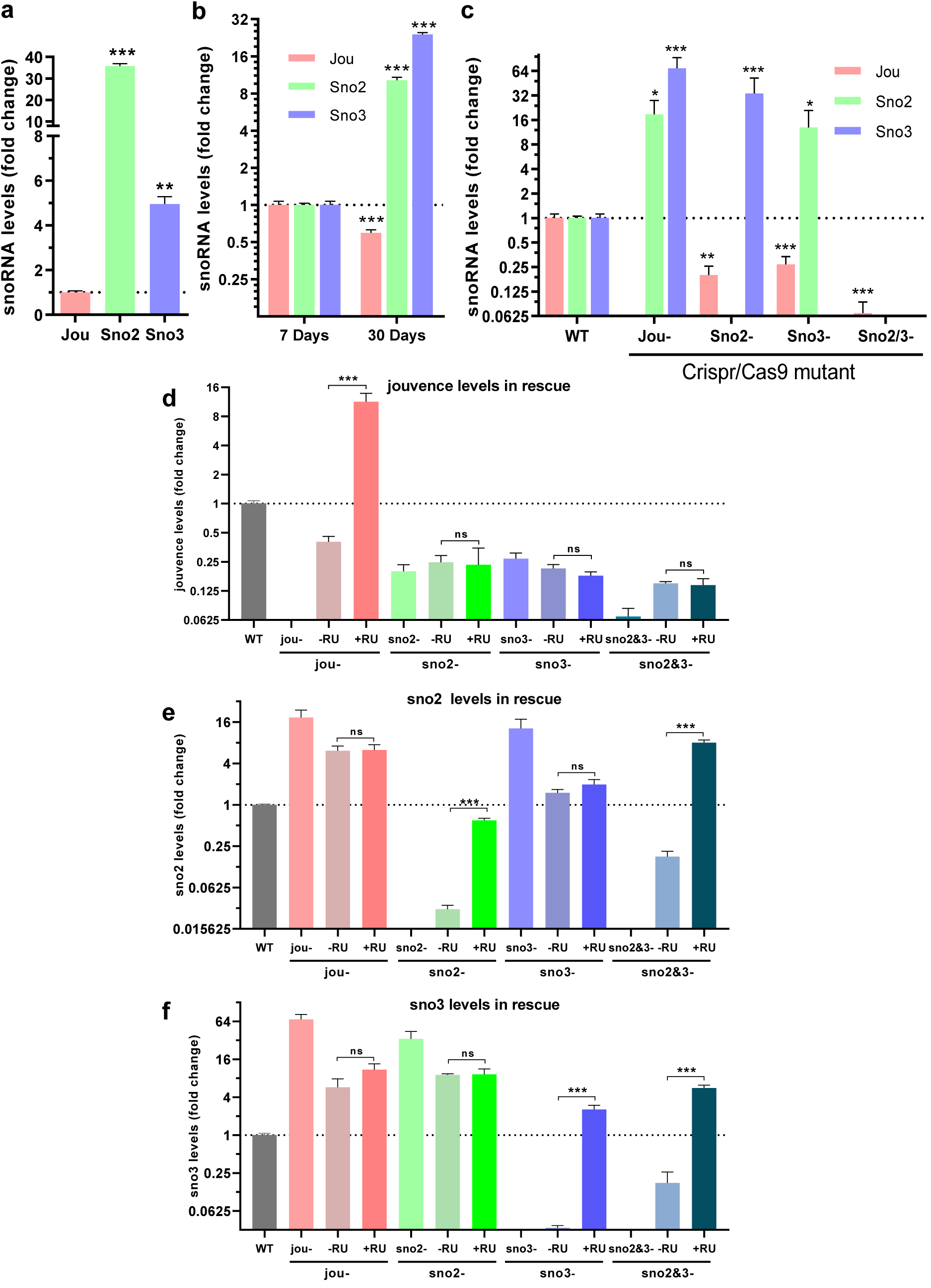
Expression levels of snoRNAs. **a)** Expression level of sno-2 and sno-3 compared to jouvence in young (7 days old) wild type flies (n=3). **b)** Expression levels of each snoRNA in wild type old flies (30 days old) compared to their respective expression in young flies (n=3). **c)** Expression levels of each snoRNA in each CRISPR/Cas9 mutant compared to their respective expression in wild type flies. **d-e-f)** Expression levels of each snoRNA in each CRISPR/Cas9 mutant and their rescue using the Mex-GS driver (Mex-GS,Crispant > UAS-snoRNA, + and – RU). **d)** jou level, **e)** sno2 level, **f)** sno3 level. Statistics: *p < 0,05; **p < 0,005; ***p < 0,0005. Errors bars represent the mean ± S.E.M. (p-value were calculated on the ΔΔCt using the two-tailed using the two-tailed unpaired T-test).

### Each snoRNA pseudouridylates a specific ribosomal-RNA site

In the previous study, we demonstrated that *jou* snoRNA pseudouridylates the *D. melanogaster (Dm)* rRNA at the positions 18S-1426 and 28S-1178 (Soulé et al., 2020). To get more insights into the molecular functions of snoRNAs present in the same cluster: sno-3 and sno-2, we first verified if each of them indeed targets the rRNA, as expected based on their complementarity to rRNA sequence (Kiss, 2020; Ye, 2007; Charette and Gray, 2000; Ofengand and Bakin, 2017). To this goal we performed a HydraPsiSeq analysis, a quantitative Psi mapping technique that relies on specific protection observed for pseudouridine residues upon hydrazine/aniline cleavage (Marchand et al., 2020). Analysis of HydraPsiSeq score values for all potentially modified *Dm* rRNA pseudouridytaion positions, allowed detection of 5 sites showing substantial and reproducible change of protection in F4-seleted cells, compared to Wild-Type control (Canton-S) (Figure 3a,b). Two of the affected positions correspond to already characterized jou targets (18S-1426 and 28S-1178), while 3 other locations (18S-1111, 28S-1170 and 28S-3085) were not yet attributed to known snoRNAs (Ofengand and Bakin, 1997; Song et al., 2023).

**Figure 3.**
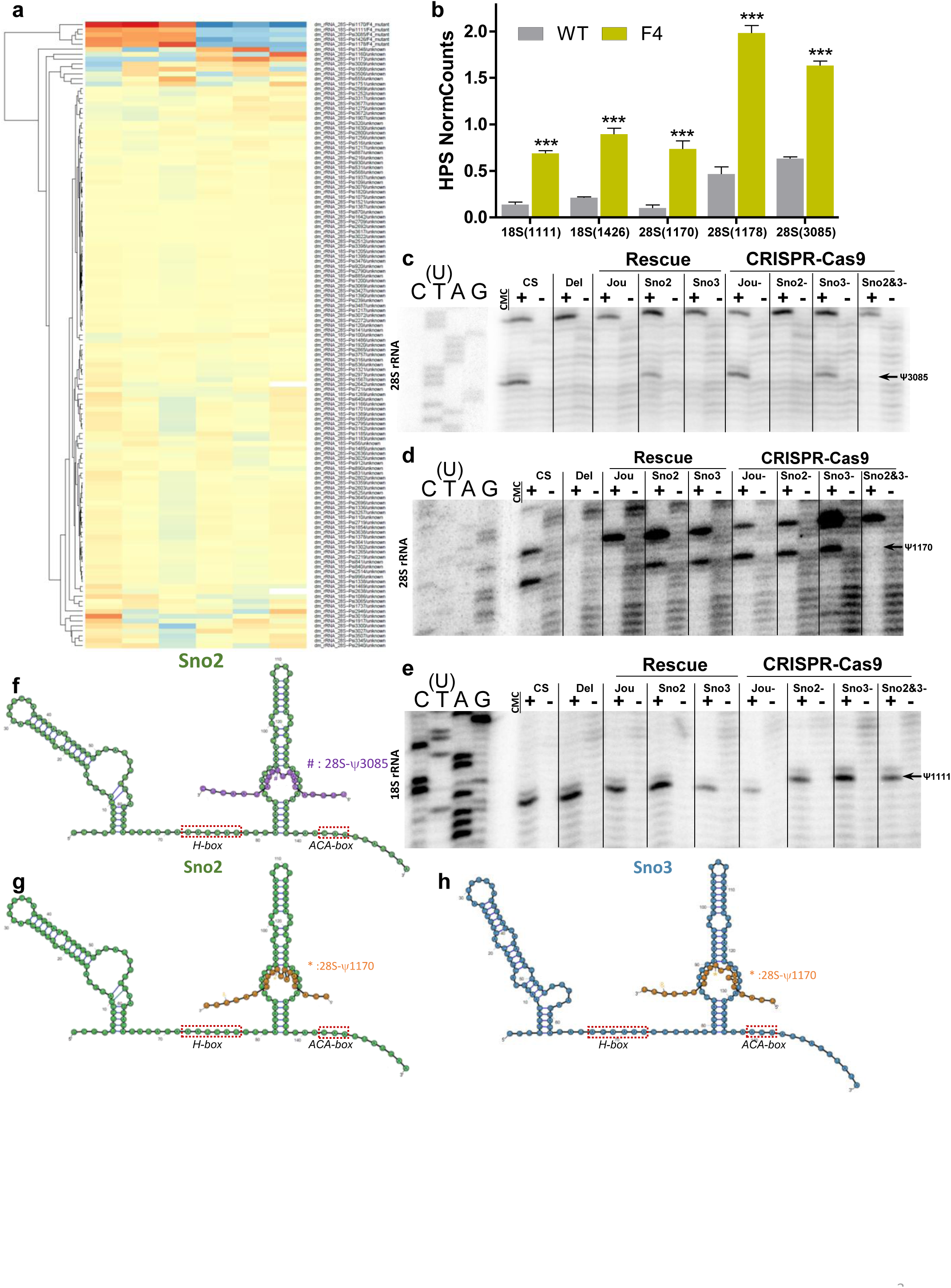
Pseudouridylation’s sites directed by each snoRNAs of the cluster. a) HydraPsiSeq heat-map representing the differential amount of pseudouridylation on rRNA sites. The first triplicates are wild type, the next triplicate are F4-deletion which includes the 3 snoRNAs, and on the right is the position of the pseudouridylation on the rRNA. **b)** HydraPsiSeq (HPS) normalized counts representing the differential pseudourydilation between wild type (WT) and F4 on the five identified significant sites. Statistics: ***p < 0,0005. Errors bars represent the mean ± S.E.M. (p-value were calculated using the two-tailed unpaired T-test). **c)** The sno2 pseudouridylates the 28S-rRNA at position 3085 in WT controls, which is missing in F4-deletion, and rescued in gut targeted sno2 (Myo1A-Gal4>sno2). This pseudouridylation is also missing in flies CRISPR/Cas9 mutated for sno2 and sno2&3. **d)** The sno2 and sno3 pseudouridylates the 28S-rRNA at position 1170 in WT controls, which it is missing in F4-deletion, and rescued in gut targeted sno2 or sno3 (Myo1A-Gal4>sno2 or sno3). This pseudouridylation is also missing in flies CRISPR/Cas9 mutated for sno2&3. **e)** The pseudouridylation on the 18S-rRNA at position 1111 is not directed by any of the snoRNA of the cluster, since this pseudouridylation is present in the F4-deletion, and in each CRISPR/Cas9 mutants. **f)** Putative interaction between sno2 and its targeted pseudouridylation site 28S-Ψ - 3085. **g)** Putative interaction between sno2 and its targeted pseudouridylation site 28S-Ψ-1170. **h)** Putative interaction between sno3 and its targeted pseudouridylation site 28S-Ψ-1170.

Complementary analysis of rRNA pseudouridylation pattern was performed by CMCT-RT-based approach. In brief, RNA is treated by CMCT reagent modifying U and Psi residues and CMC covalent adduct at Us is removed by alkaline treatment. The residual RT stops observed on the denaturing PAGE correspond to Psi residues. The results of this analysis demonstrated that Psi at position 28S-3085 is guided solely by sno-2 (Figure 3c), since the RT stop is missing in sno-2 CRISPR/Cas9 mutant and efficiently requested by the targetted expression of UAS-sno-2. Modification at the position 28S-1170 is guided by either sno-2 and/or sno-3 (Figure 3d), since this residue is missing only in doubly deleted sno-2&3 CRISPR/Cas9 mutant, but not in sno-2 and sno-3 mutants, and targetted expression of UAS-sno-2, or UAS-sno-3, which allows to restore modification (Figure 3d). Surprisingly, in contrast to HydraPsiSeq data, we were not able to detect a missed pseudo-U at the position 18S-1111, neither in F4-deletion, neither in any CRISPR/Cas9 mutants (Figure 3e). Although we have not yet the precise explanation for this discrepancy, we suspect that this could be due to a slight polymorphism at this base. Figure 3f,g,h schematically shows the putative 2D-structure of the sno2 and sno3, and their complementarity to detected pseudouridylation sites in rRNA.

Altogether, these experiments confirm that each of the three snoRNAs present in *jou* cluster guides pseudouridylation of the rRNAs at a specific site, indicating that these snoRNAs are indeed a *bona fide* H/ACA guides, and that our CRISPR/Cas9 replacement mutations lead to a non-functional snoRNAs.

### The three snoRNAs affects the amount of ribosomes and translational efficacy

Since the snoRNAs, and particularly the H/ACA boxes, are known to affect the biogenesis and the stability of the ribosomes (Kiss, 2002; Ye, 2007), first, to asses if the biogenesis or the stability of ribosomes is affected, we have determined the amount of the ribosomes by targetting the expression of the RPL10-GFP (Huang et al., 2013; You et al., 2021) in the enterocytes of the gut (F4,Myo-Gal4>UAS-RPL10-GFP). The quantification of the GFP-fluorescence reveals that the F4-deleted flies present about only half of the amount of ribosomes compared to the Wild-Type control (Figure 4a,b,i), while the targeted re-expression of each snoRNAs improves and even rescues the amount of ribosomes (Figure 4c,d,e,f,g,h,i). Second, since the snoRNAs have also been shown to affect the ribosomal ligand binding and the translation (Jack et al., 2011), we performed a puromycin labelling experiment (Haussmann et al., 2022 ; Schmidt et al., 2009) in which the puromycin, an analog of the tyrosine, is incorporated within the nascent peptide chain, and thus allows to quantify the protein synthesis rate. The F4-deleted flies present a strong decrease of protein synthesis rate, as well as each CRISPR/Cas9 mutants (Figure 4j-u), while the targeted re-expression of each snoRNAs within the enterocytes of the gut rescues it. Finally, to more directly investigate the translation efficacy of mRNA, we perform a TRAP (Translational Ribosomal Affinity Purification) assay (Heiman et al., 2014), by specifically targeting the ribosomal protein (RPL13) tagged with a Flag (RLP13A-FLAG) (Huang et al., 2019), in the enterocytes using the Myo1A-Gal4 driver, in the F4-deleted flies. Following the TRAP, we perform a RT-qPCR on specific selected genes involved principally in metabolic pathways. The translation level is importantly reduced in F4-deleted flies compared to Wild-Type Control for the genes ACC, Lip4, and FOXO (Figure 4v,w,x). In a more global overview, extending our study to some other genes, as FASN1, FA2H, NPC1 and NPC2, we remark that several of them show a tendency to be decreased (Figure 4z), although these last are not individually statistically different. Interestingly, we also notice that not all the investigated genes are decreased, as for instance the gene ninaD (Figure 4y), which the expression level is importantly increased in F4-deleted flies (Soulé and Martin, 2020), allowing to rule-out that this effect is likely not solely due to a decrease in the general amount of ribosomes. Altogether, these three independent approaches (GFP-Fluorescence, Puromycin, and TRAP) indicate that both the amount of ribosomes, as well as the translation efficacy are reduced by the mutations of these snoRNAs.

**Figure 4.**
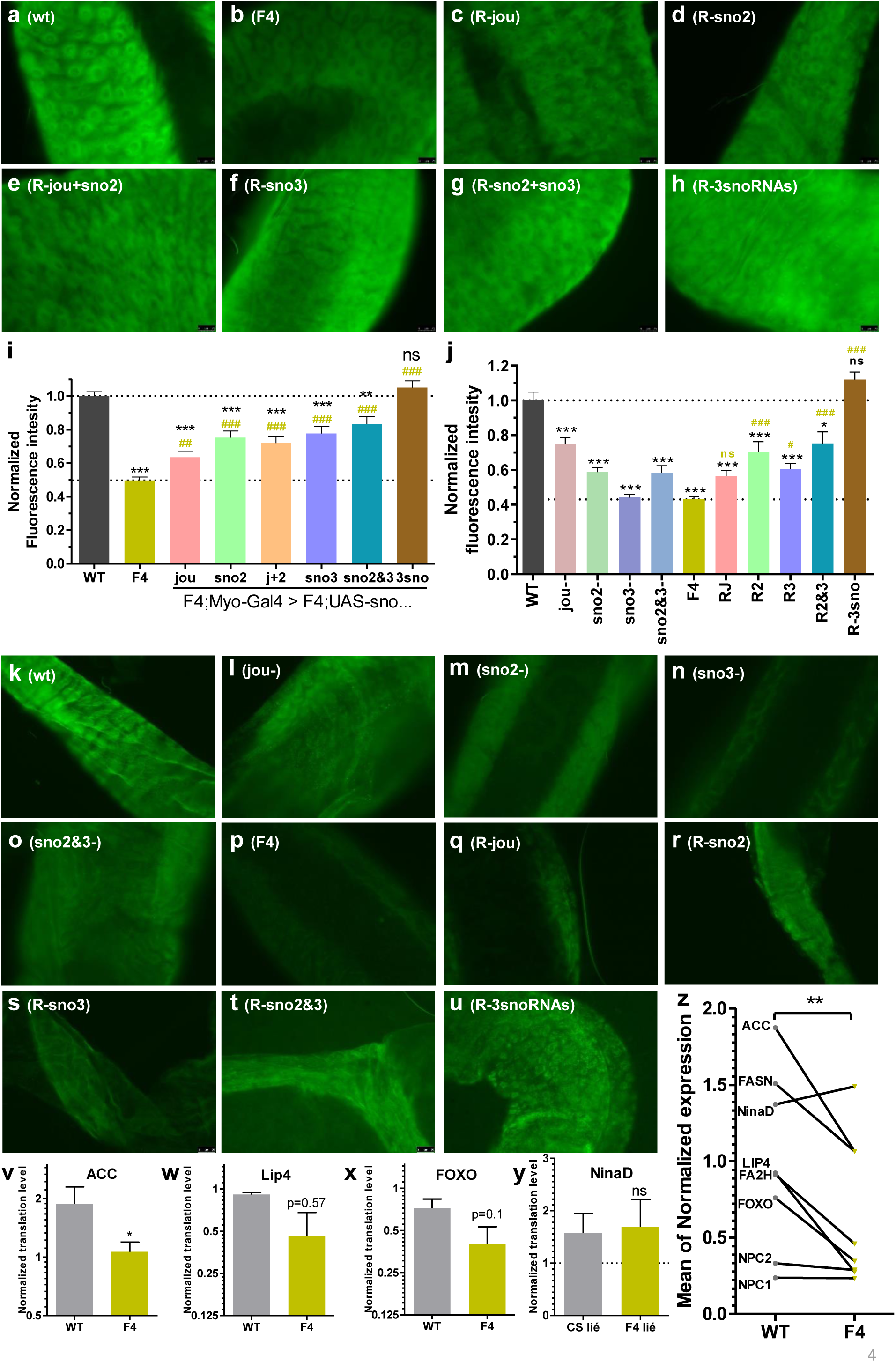
SnoRNAs affect ribosome regulation and protein synthesis rate. **a-h)** RPL10-GFP imaging in enterocytes of mid-gut in wild type, mutants and rescues (F4,Myo-Gal4 > F4; UAS-sno, UAS-RPL10-GFP). **a)** in wild-type flies ; **b)** in F4 flies; **c)** in F4 flies rescued with jouvence; **d)** in F4 flies rescued with sno2; **e)** in F4 flies rescued with jouvence and sno2; **f)** in F4 flies rescued with sno3; **g)** in F4 flies rescued with sno2 and sno3; **h)** in F4 flies rescued with the three snoRNA (jouvence + sno2 + sno3). **i)** Ribosome quantification by normalized fluorescence intensity of RPL10-GFP in wild-type, F4, and rescue of every snoRNA in enterocytes. WT: n=27, F4: n=20, rescue-jou: n=27, rescue-sno2: n=29, rescue-jou+sno2: n=25, rescue-sno3: n=28, rescue-sno2+sno3: n= 30, rescue-3sno: n=21. **j)** Protein synthesis rate quantified by puromycin-assay. Assay on flies mutated by CRISPR/Cas9 for jouvence, sno2, sno3, or sno2&3, or for the deletion of each snoRNAs (F4) and on F4 flies rescued with jouvence (RJ), sno2 (R2), sno3 (R3), sno2&3 (R2&3), or all three snoRNAs (R-3sno). **i,j)** Statistics: *p < 0,05; **p < 0,005; ***p < 0,0005 compared to wild type; #p < 0,05; ##p < 0,005; ###p < 0,0005 compared to F4. Errors bars represent the mean ± S.E.M. (p-value were calculated using one-way ANOVA with Turkey post-tests). **k-u)** Puromycine-assay imaging in enterocytes of mid-gut in wild type, mutants and rescue (F4,Myo-Gal4 > F4;UAS-sno) ; **k)** in wild-type flies ; **l)** in Crispant-jou; **m)** in Crispant-sno2; **n)** in Crispant-sno3; **o)** in Crispant-sno2&3; **p)** in F4 flies ; **q)** in F4 flies rescued with jouvence; **r)** in F4 flies rescued with sno2; **s)** in F4 flies rescued with sno3; **t)** in F4 flies rescued with sno2 and sno3; **u)** in F4 flies rescued with the three snoRNA (jouvence + sno2 + sno3). **v-x)** Normalized translation levels quantified by Translational Ribosomal Affinity Purification (TRAP) representing the translatability of the mRNA normalized by total mRNA compared between wild type and F4. **v)** for ACC mRNA ; **w)** for Lip4 mRNA ; **x)** for FOXO mRNA, **y)** for ninaD mRNA. **z)** Mean of normalized expression of few other genes, showing that although these last genes are not staticically different, altogether they show a common decrease between wild type (WT) and F4 deletion. Statistics: *p < 0,05 compared to wild-type. Errors bars represent the mean ± S.E.M. (p-value were calculated on the ΔΔCt using the one-tailed unpaired T-test with a decrease assumption).

### The specific mutation of each snoRNA deregulates metabolic parameters

To assess the physiological consequences of the deregulation of the ribosomal machinery, and since several metabolic pathways genes are deregulated in F4-deleted flies as revealed by the previous RNA-Seq performed on the gut (Soulé et al., 2020 ; Al Issa et al., 2026), we quantify the level of triglycerides (TG) and sterols (including the cholesterol). Similar to the previously reported for the F4-deleted flies (Soulé and Martin, 2020; Al Issa et al., 2026), the CRISPR/Cas9 mutation of each snoRNA increases the triglycerides level (Figure 5a,c). Interestingly, this effect correlates with the expression level of FASN1 and ACC genes, the two main limiting genes involved in de-novo fatty-acid synthesis (Heier and Kühnlein, 2018), whose are decreased in jou, sno2, and sno2&3 CRISPR/Cas9 mutants, but increased in sno-3 mutant (Figure 5d,g). Moreover, the expression level of these two genes is also rescued by the re-expression of the sno2 or sno3 within the enterocytes of the gut (Myo1A-Gal4>UAS-sno) placed in the corresponding Crispant-mutant: FASN1 by sno2 and sno3 (Figure 5e,f), whereas ACC by sno2 only (Figure 5h), while jou did not rescue any of them. Interestingly, in parallel, the triglycerides level is also rescued (or at least partly) in a similar manner: partly rescued by sno2, well rescued by sno3, and not rescue by jou (Figure 5b).

**Figure 5.**
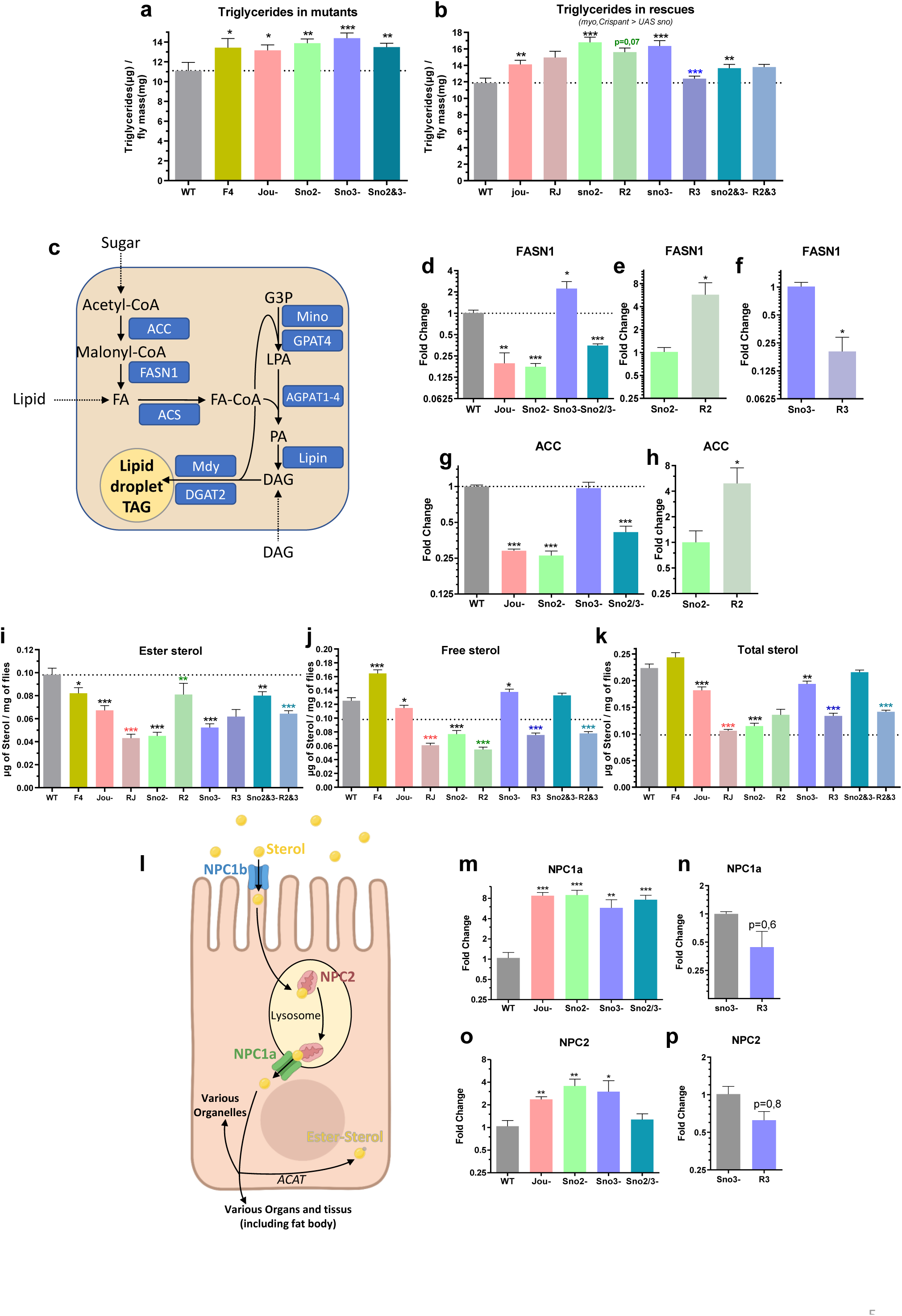
SnoRNAs impact on lipid metabolism. **a)** Triglycerides levels in wild-type, F4, and CRISPR/Cas9 mutants. **b)** Triglycerides levels in CRISPR/Cas9 mutant flies rescued for the corresponding snoRNA using Myo-Gal4>UAS-sno… in corresponding Crispant mutant). **a,b)** Quantification on whole 7 day-old fly extract. All mutants are either compared to the wild-type for the corresponding triglycerides (*in black) or to the corresponding snoRNA mutant in color (red to jou-, green to sno2-, dark-blue to sno3- , cyan to sno2&3-). p-value were calculated using one tailed t-test assuming increase in triglyceride for mutant and decrease for rescue. **c)** Schematic drawing summarizing the main genes involved in triglycerides synthesis. **d-h)** Quantification on dissected gut of 7 day-old flies. **d)** FASN1 expression levels in wild-type and CRISPR/Cas9 mutants. **e)** FASN1 level is rescued by sno2 driven by Myo-Gal4 (Myo-Gal4>UAS-sno2 in Crispant-sno2: R2). **f)** FASN1 level is rescued by sno3 driven by Myo-Gal4 (Myo-Gal4>UAS-sno3 in Crispant-sno3: R3). **g)** ACC expression levels in wild-type and CRISPR/Cas9 mutants. **h)** ACC level is rescued by sno2 driven by Myo-Gal4 (Myo-Gal4>UAS-sno2 in Crispant-sno2: R2). **i,j,k)** Quantification on whole 7 day-old fly extract. Ester-sterol, free-sterol and total-sterol levels in wild-type and CRISPR/Cas9 mutants, and in flies rescued for the corresponding snoRNA using Myo-Gal4>UAS-sno… in corresponding Crispant mutant. All mutants are either compared to the wild-type for the corresponding sterol (* in black) or to the corresponding snoRNA mutant in color (red to jou-, green to sno2-, dark-blue to sno3-, cyan to sno2&3-) for the rescue. Errors bars represent the mean ± S.E.M. (p-value were calculated using two-tailed unpaired T-test). **l)** Schematic drawing summarizing the main genes involved in sterol absorption and metabolism. **m-p)** Quantification on dissected gut of 7 day-old flies. **m)** NPC1a expression levels in wild-type and CRISPR/Cas9 mutants. **n)** NPC1a level is partially rescued by sno3 driven by Myo-Gal4 (Myo-Gal4>UAS-sno3 in Crispant-sno3: R3). **o)** NPC2 expression levels in wildtype and CRISPR/Cas9 mutants. **p)** NPC2 level is partially rescued by sno3 driven by Myo-Gal4 (Myo-Gal4>UAS-sno3 in Crispant-sno3: R3). Statistics: *p < 0,05; **p < 0,005; ***p < 0,0005. Errors bars represent the mean ± S.E.M. **d-h** and **m-p)** p-value were calculated on the ΔΔCt using the two-tailed unpaired T-test.

Similarly, we also quantify the level of sterols (including the cholesterol). As previously reported (Soulé and Martin, 2020; Al Issa et al., 2026), the F4-deleted flies show a decrease of sterol-ester, while the CRISPR/Cas9 mutation of jou, sno2, and sno3 also present a decrease of sterol-ester, and to a less extent, the sno2&3 together (Figure 5i,j,k). Interestingly, also here, the deregulation of this metabolic parameter is supported by the deregulation of some specific genes involved in sterol metabolism, as NPC1a (Voght et al., 2007; Philips et al., 2008), and NPC2 (Huang et al., 2007) (Figure 5l). Indeed, the expression level of NPC1a is increased in each CRISPR/Cas9 mutants ; jou, sno2, sno3 and sno2&3 (Figure 5m), while the NPC2 is also increased in jou, sno2 and sno3 mutants, but not in the double mutants (sno2&3) (Figure 5o). However, the expression level of these two genes is rescued only by the re-expression of the sno3 within the enterocytes of the gut (Myo1A-Gal4>UAS-sno3) in its cognate Crispant-mutation (sno3) (Figure 5n,p). In overall, altogether, the deregulation of these two physiological parameters (TG and sterol) by each CRISPR/Cas9 mutants, related to the deregulation of specific metabolism genes, attests that the deregulation of the ribosomal machinery specifically in the enterocytes of the gut leads to general physiological defects.

### The specific mutation of each snoRNA also affects the longevity

Since we have formerly reported that the deletion of the 3 snoRNAs altogether (F4-deletion) decreases the lifespan of the flies (Soulé et al., 2020), while the genetically targetted re-expression of each of them rescues, at least in part, the longevity (Al Issa et al., 2026), we wonder if the deletion of a single snoRNAs also affects the lonvevity. In contrast to the F4-deletion, the CRISPR/Cas9 mutation (also commonly named: Crispant) of a single specific snoRNA affects differently the longevity of the flies (Figure 6a). In jou-Crispant (in which the sno2 and sno3 are still present, and even increased: see Figure 2a,b,c), the longevity is increased. The longevity is also slightly increased for the sno2-Crispant (in which jou is decreased, while sno3 is increased: see Figure 2c), or sno3-Crispant (in which jou is decreased and sno2 increased: see Figure 2c). However, when the sno2&3 are mutated together (sno2&3-Crispant) (in which only jou is very weakly present, and even almost undetectable: see Figure 2c), the longevity is decreased, and interestingly, similar to F4-deletion. To complement, the targeted and specific re-expression of each snoRNA within the epithelium of the gut, in its corresponding Crispant-mutant, when the expression is induced only in adulthood, using Mex-GS, rescues the longevity of the flies (Figure 6b,c,d,e) (decreased after induction by feeding RU only in adulthood in jou, sno2 and sno3, and increased in sno2&3). In overall, altogether, these results demonstrate that each snoRNAs is involved, each in their own way, in the determination of the longevity of the flies.

**Figure 6.**
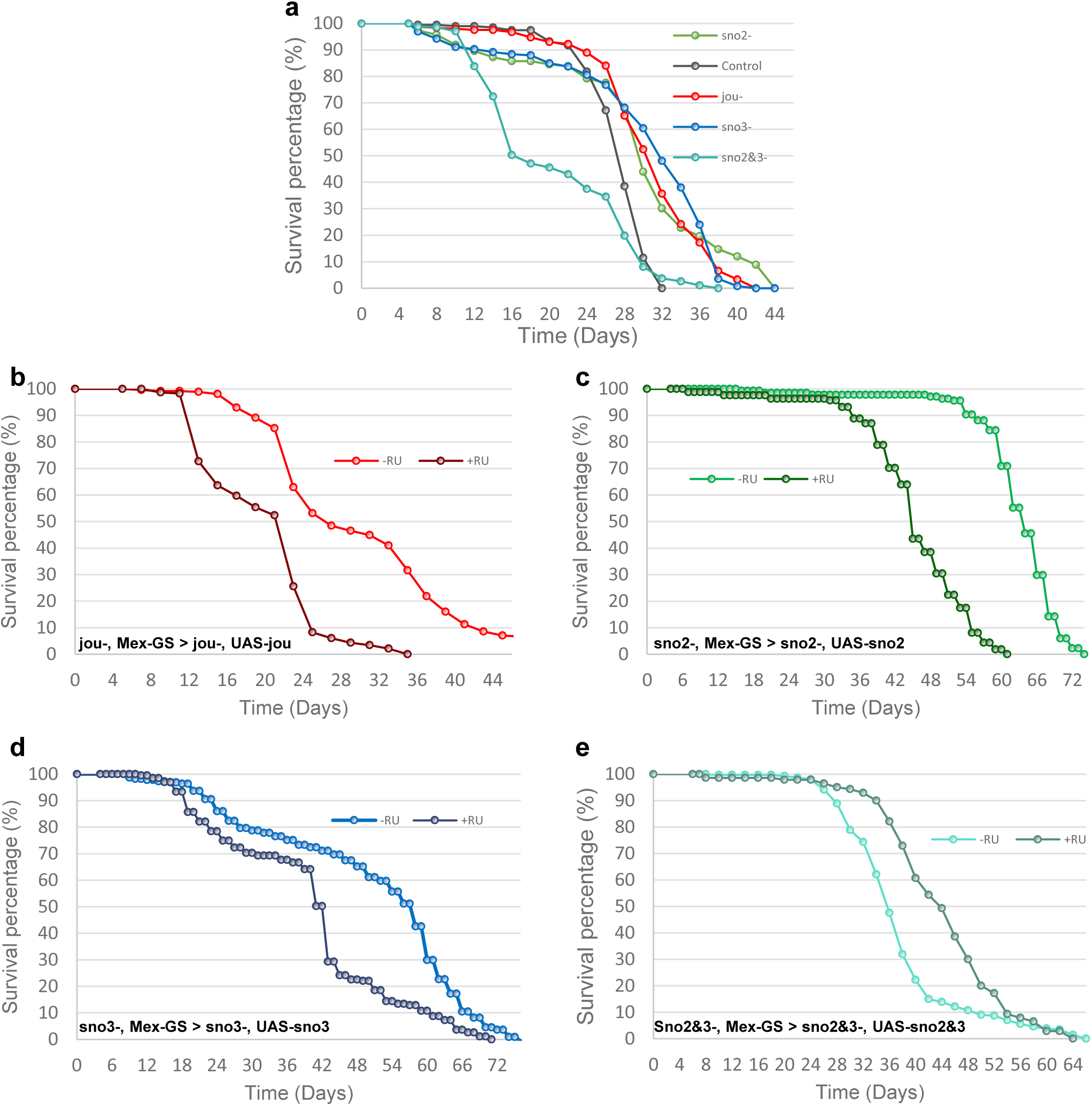
Targeted expression of each snoRNA in the enterocytes rescues the longevity. Longevity test result (survival curve – decreasing cumulative) **a)** Control flies and CRISPR/Cas9 mutant for jouvence (jou-), sno2 (sno2-), sno3 (sno3-), and sno2&3 (sno2&3-) (p-value<0,0001 for each mutant compared to the control). **b)** jou-,Mex-GS > UAS-jou, flies fed with or without RU486 (p-value<0,0001). **c)** sno2-,Mex-GS > UAS-sno2-, flies fed with or without RU486 (p-value < 0,0001). **d)** sno3-,Mex-GS>UAS-sno3-, flies fed with or without RU486 (p-value < 0,0001). **e)** sno2&3-,Mex-GS > UAS-sno2&3-, flies fed with or without RU486 (p-value < 0,0001). Log-rank test p-values calculated by OASIS Software (Yang et al., 2011).

### The specific mutation of each snoRNA leads to neurodegeneration in old flies

Since each snoRNA affects the longevity of the flies, and more specifically increase it, we wonder if this increase of longevity is also correlated to modification in the brain of the flies. In other words, do the flies that live longer have less neurodegenerative lesionsSimilar to the previously reported for the F4-deleted flies (deletion of the 3 snoRNAs altogether) (Soulé and Martin, 2020; Al Issa et al., 2026), the CRISPR/Cas9 mutation of each specific snoRNA also leads to more neurodegenerative lesions in old flies (Figure 7a-f,p), compared to control Wild-Type flies. Moreover, the targeted re-expression of each snoRNA specifically within the epithelium of the gut (Mex-GS>UAS-sno…) induced by feeding RU only in adulthood, in each corrresponding Crispant-mutant, differently rescues the number of neurodegenerative lesions (Figure 7g-o,q). In particular, jou do not rescue, whereas sno2 rescues perfectly, while sno3 or sno2&3 only partly rescue the number of lesions. Interestingly, in overall, these last results correlate, at least in their major part, to the rescue of the metabolic parameters, as triglycerides and sterols (Figure 5), but intriguingly, not to the longevity for jou, sno2, and sno3, but perfectly for the sno2&3 (Figure 6).

**Figure 7.**
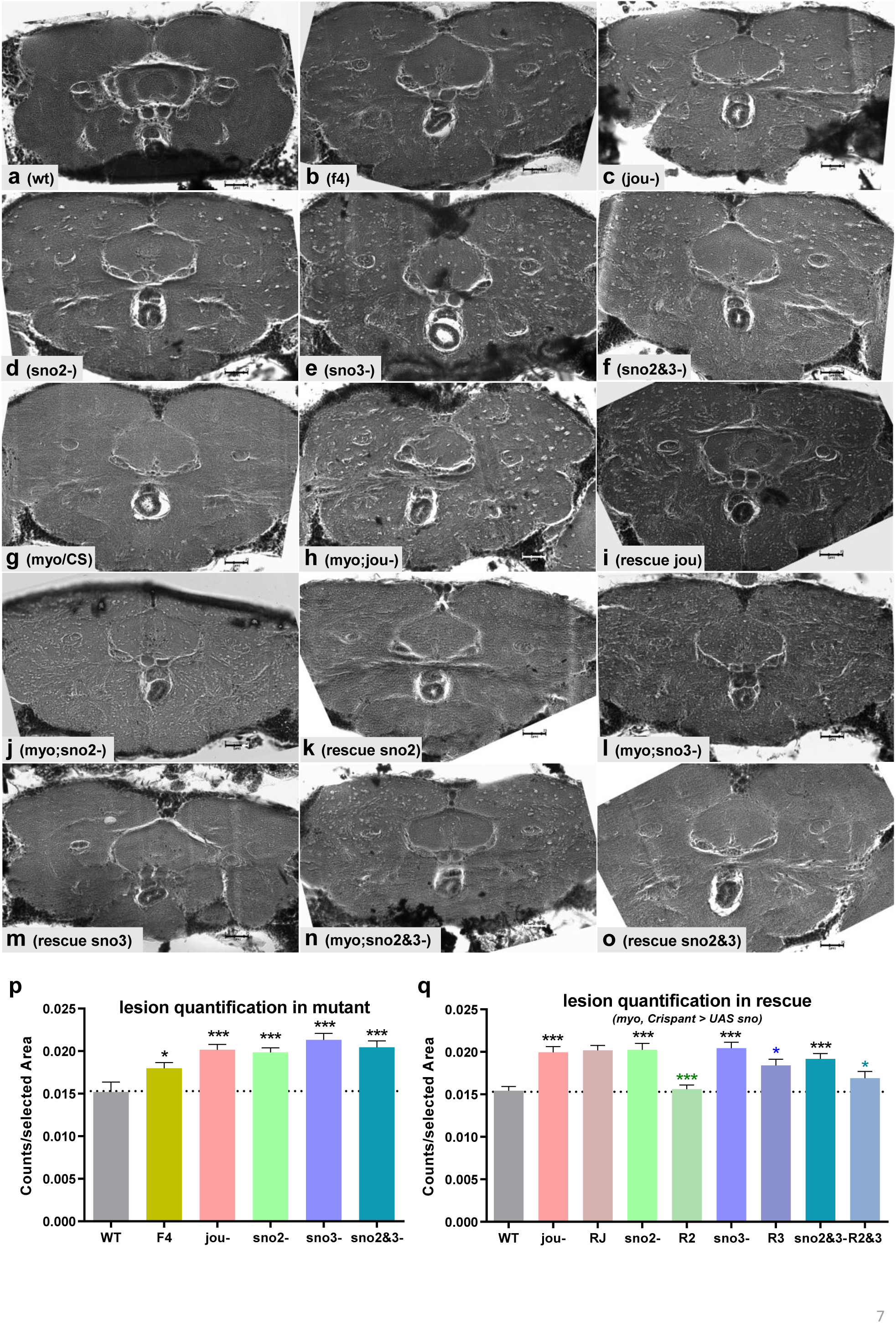
SnoRNAs protect against neurodegeneration. **a-o)** Microphotography showing the vacuoles (lesions) in old flies in **a)** Wild-Type ; **b)** F4 mutant/deletion for the 3 snoRNAs ; **c)** CRISPR/Cas9 mutant jouvence ; **d)** CRISPR/Cas9 mutant sno2 ; **e)** CRISPR/Cas9 mutant sno3 ; **f)** CRISPR/Cas9 mutant sno2&3 ; **g-o)** rescue (Myo-Gal4 > UAS-sno…. in each corresponding Crispant-mutant). G) Myo-Gal4/CS (control); **h)** Myo-Gal4,jou- (control/mutant)) ; **i)** rescue jouvence : Myo-Gal4,jou- > UAS-jou ; **j)** Myo-Gal4,sno2- (control/mutant) ; **k)** rescue sno2 : Myo-Gal4,sno2->UAS-sno2 ; **l)** Myo-Gal4;sno3-(control/mutant); **m)** rescue sno3: Myo-Gal4,sno3->UAS-sno3; **n)** Myo-Gal4,sno2&3-(control/mutant) ; **o)** rescue sno2&3: Myo-Gal4,sno2&3->UAS-sno2&3; **p)** Quantification of neurodegenerative lesions in Wild-Type and different mutants compared to CS (Wild-Type). CS: n=22 ; F4 : n=46 ; jou- : n=46 ; sno2- : n=52 ; sno3- : n= 43 ; sno2&3- : n=42. **q)** Quantification of neurodegenerative lesions in mutant flies rescued with their corresponding snoRNAs compared to Myo-Gal4/CS (* in black), and to the corresponding control mutants in color. Myo-Gal4/CS: n=54 ; Myo-Gal4,jou- : n=42 ; R = rescue: RJ: n=35 ; Myo-Gal4,sno2- : n=44 ; R2 : n= 55 ; Myo-Gal4,sno3- : n=48 ; R3 : n= 50 ; Myo-Gal4,sno2&3- : n=51 ; R2&3 : n= 46. Statistics: *p < 0,05; **p < 0,005; ***p < 0,0005 compared to Wild-Type. Errors bars represent the mean ± S.E.M. (p-value were calculated using one tailed t-test assuming increase in lesions for mutants and decrease for rescue).

## Discussion

We have demonstrated here that each snoRNAs of the small cluster of 3 snoRNAs including *jouvence* is involved, each in their own way, in the lifespan determination, metabolic parameters, and neurodegeneration/neuroprotection. In the continuity of our characterisation of a small cluster of 3 snoRNAs, by generating specific CRISPR/Cas9 mutation of each of them, first, using two independant approaches (HydraPsiSeq and CMCT-RT analysis of pseudouridylation), we have demonstrated, at molecular level, that each of them pseudouridylates a specific site on ribosomal RNAs, suggesting a higher complexity in rRNAs pseudouridylation than previously expected. In a step further, to temptingly decipher the consequence of the pseudouridylation on the ribosomal machinery and functionality, using two *in-vivo* independent approaches, we showed that each snoRNA mutation affects the amount of ribosomes as well as the translational efficacy. Indeed, first, targeting a GFP-tagged ribosomal proteins (RPL13-GFP), reveals a decrease in the amount of ribosomes, suggesting that the pseudouridylation either affects the primary ribogenesis, or more likely, affects their stability. Further experiments will be required to temptingly discern between these two hypothesis (Jack et al., 2011). Second, the puromycin incorporation has revealed that the mutation of each snoRNA reduces the level of general translation, a result in accordance with a previous report (Jack et al., 2011). Finally, the TRAP experiment, based on the targeted Flag-tagged ribosomal protein (RpL13A-Flag) (Huang et al., 2019) specifically in the enterocytes, has allowed to reveal that the translation of some specific genes is affected; some being reduced, while some others are not affected, indicating that site-specific pseudouridylation specifically affects the translation of some genes, but not others.

In an attempt to link this molecular defects to more general organismal phenotypes, we have demonstrated that some genes involved specifically in the lipid metabolism, as FASN1 and ACC, two limiting genes of fatty-acid synthesis pathway (Heier and Kühnlein, 2018), are deregulated in CRISPR/Cas9 mutants, while they are rescued by the targetted re-expression of their corresponding snoRNA, which consequently, correlate with the level of triglycerides. Similarly, two genes involved in sterol metabolism (NPC1a and NPC2), also correlates with the level of sterol-ester, reinforcing also here, the link betwen the expression of the snoRNAs within the gut, the expression of these specific metabolic genes, and the metabolic parameters. It goes without saying that a fine regulation of the metabolism all along the life is a prerequisite for the healtspan. Thus, even a tiny chronic deregulation of some metabolic parameters, as triglycerides and sterol, including the cholesterol, could affect, in long term, the healthy lifespan (Broughton et al. 2005; Al Issa et al., 2026). Here, the longevity is affected in each CRISPR/Cas9 mutants (increased in Crispant-jou, Crispant-sno2 and Crispant-sno3, but decreased in double Crispant-sno2&3). More importantly, it is rescued, at least partially, by the re-expression of the each snoRNA within the enterocytes. However and astonishingly, in contrast to the deletion of the 3 snoRNAs (F4-deletion) which decreases the longevity (Soulé et al., 2020 ; Al Issa et al., 2026), the mutation, by a non-fonctional subtitution sequence, of a single snoRNA increases the longevity. This discrepancy is likely due to the fact that the mutation of a single snoRNAs importantly affects the level of the two others (Figure 2), leading to a more complex effect on longevity than the single mutation itself, and thus complicating the interpretation of the results. Another explanation could be that the genetic targeted rescue system used, based on the P[Gene-Swith], does not always perfectly rescue the expression level of the transgene (UAS-snoRNA), and even have the general tendency to express much more than the physiological level (in other words, sometime ressembling rather to an overexpression). To resume, each of the snoRNA are required in the enterocytes of the epithelium of the gut to specifically pseudourydylates a precise and given rRNA site (Figure 8). This pseudouridylation affects the ribosomal machinery, both the amount of ribosomes, as well as the protein synthesis rate of specific genes, and more particularly the expression of the genes involved in lipids and sterol metabolism, as revealed by the transcriptomic analysis (Soulé et al., 2020). Consequently, this chronic deregulation of metabolic parameters all along the life yields to a perturbation of lipids and sterols within the pericerebral fat body and the brain (Al Issa et al., 2026), which *in fine*, perturbs the turnover if these essential cellular components, leading to neurodegenerative lesions in old flies.

**Figure 8.**
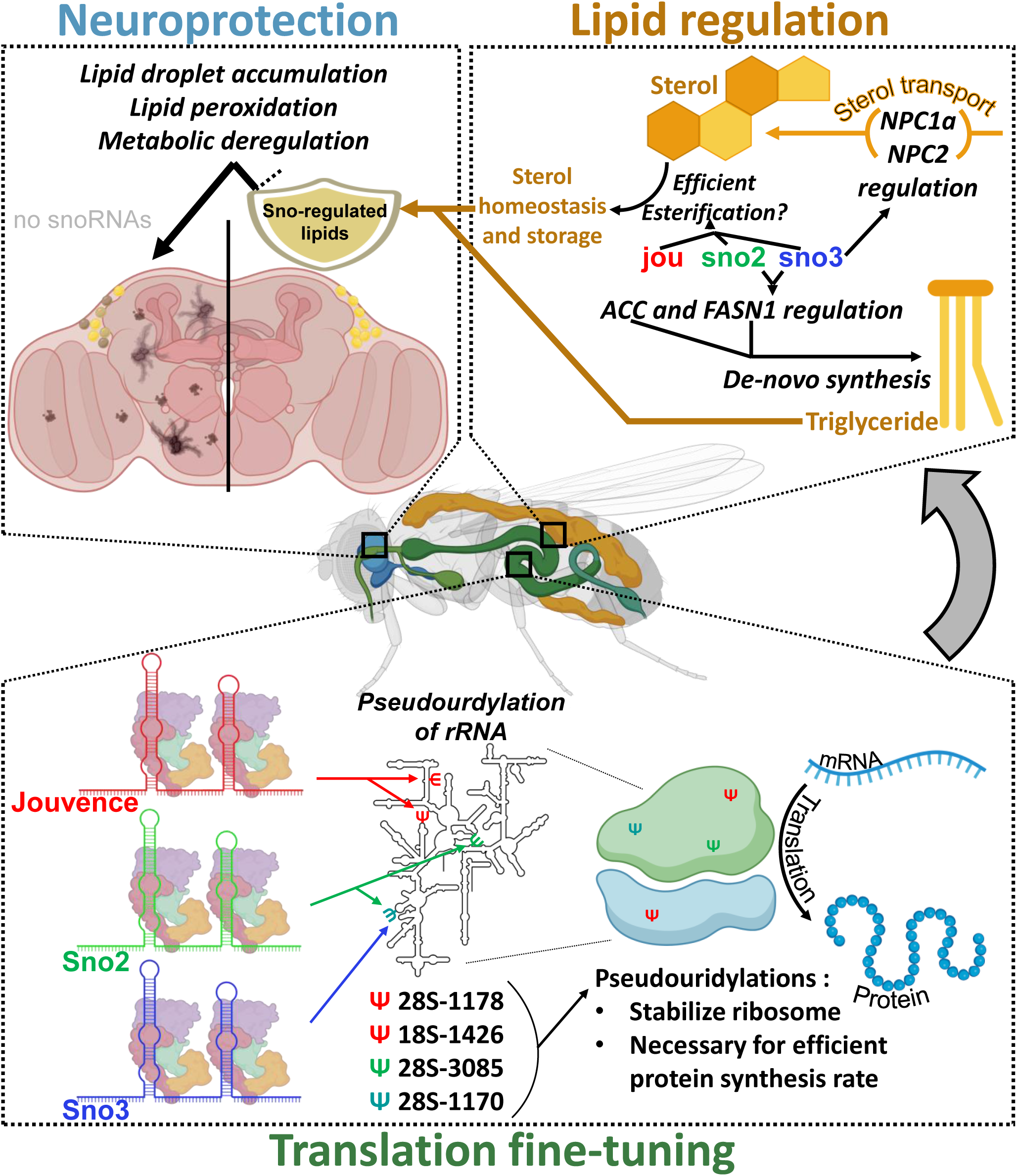
Graphical summary: Pseudouridylations of rRNA by jou, sno2 and sno3 affect ribosomal machinery and consequently neurodegeneration through lipid metabolism deregulation. Jouvence, sno2 and sno3 are responsible for four pseudouridylations sites (28SΨ1178 [jou], 18SΨ1426 [jou], 28SΨ1170 [sno2 and sno3], 28SΨ3085 [sno2]). The presence of those pseudouridylations might stabilise the ribosomes, and required for an efficient protein synthesis rate. The pseudouridylation caused by sno2 and sno3 will contribute to the regulation of triglycerides through *de-novo* synthesis (ACC & FASN1). The presense of the pseudouridylations by the 3 snoRNAs is required for efficient sterol transport and consequently sterol homeostasis. The aforementioned lipid homeostasis protects the brain against the lipid accumulation and peroxidation, protecting against accumulation of neurodegenerative lesions.

## Materials & Methods

### Drosophila lines

*Drosophila melanogaster* flies were grown on standard medium (1.04% agar, 8% cornmeal, 8% brewer yeast and 2% nipagin as a mold inhibitor) at 25°C, 12:12 light:dark cycle in a humidity-controlled incubator. For ageing experiments, 15 adult female flies were crossed with 10 males per vial, and were transferred to new food vials every 2 days. Wild-Type *Canton-S* (CS) flies were used as control. The results described in this study were obtained from females. All genotypes were outcrossed to the wild-type CS (cantonization) at least 6 times to homogenize the genetic background. RU486 (Mifepristone) (*Sigma-Aldrich*, Cat. #M8046) to induce the Gene-Switch activity was dissolved in ethanol and mixed into the media when preparing food vials. RU486 doses used were 25 µg.mL^-1^ final concentration. Myo1A-Gal4 was kindly provided by B.A. Edgar (Heidelberg, Germany), CG8997-GS from H. Tricoire (Paris, France). Mex-GS (in VK02 site), UAS-jou (inserted randomly on the third chromosome), UAS-sno2 (in VK27 site), UAS-sno3 (in VK27 site), UAS-sno2&3 (in VK27 site), and UAS-3snoRNAs (in attP2 site), have been generated in our laboratory (Soulé et al., 2020). These various transgenic lines were then introduced into their corresponding CRISPR/Cas9 mutant genetic background by standard genetic crosses.

### CRISPR/Cas9 mutants genesis

The CRISPR/Cas9 mutant of each snoRNA (jou, sno-2, sno-3 and sno-2&3 together) have been generated in subcontracting to WellGenetics (Taiwan). Briefly, in the goal to minimally affect the overall structure of the whole locus, we have opted for a replacement sequence emcompassing the H-box, which integrity is known to be esssential to the functionality of the snoRNA (Kiss, 2020). For jou, we use a 24 bp replacement, while for the sno-2 and sno-3, since these two last have a high amount of homology, we use a 55 bp replacement. Finally, for the sno-2&3, we use the same sequence as for the sno-2 and sno-3 respectiveley, to replace and mute the two snoRNAs together (Figure 1b).

### Lifespan analysis

Following amplification, flies were harvested every day after hatching. Female flies were maintained with males in fresh food vials for 4 days at a density of 25 individuals per vial. On day 4, males were removed, while mated females were placed in a cage (about 200 to 300 females per cage), as in Soulé et al. (2020). The ageing animals were transferred to fresh food every two days, and the number of dead flies was scored. The lifespan plots (survivorship curves) were generated by calculating the percentage of survivors every two days and plotting viability as a function of time (days) using log-rank test (Yang, et al. 2011).

### Quantitative RT-qPCR

Expression level of each snoRNA was measured on dissected gut or whole flies from 7-days-old female flies, as in Soulé et al. (2020). Quantitative RT-qPCR was performed on a QuantStudio-3 instrument (Applied Biosystem/ThermoFisher). For the snoRNA, we use a Taqman probe for each snoRNA (ThermoFisher) (jou = 4331348-AICSXCO; sno2= 4331348-APYMJZP; sno3= 4331348-APZTEKM), while for the encoding genes, we used the SybGreen. All assays were done in triplicate. Data were analyzed according to the ΔΔCt method, and normalized to RPL32 (ex-rp49) levels. For the encoding genes, primers used are summarized in the Table 1.

**Table 1.**
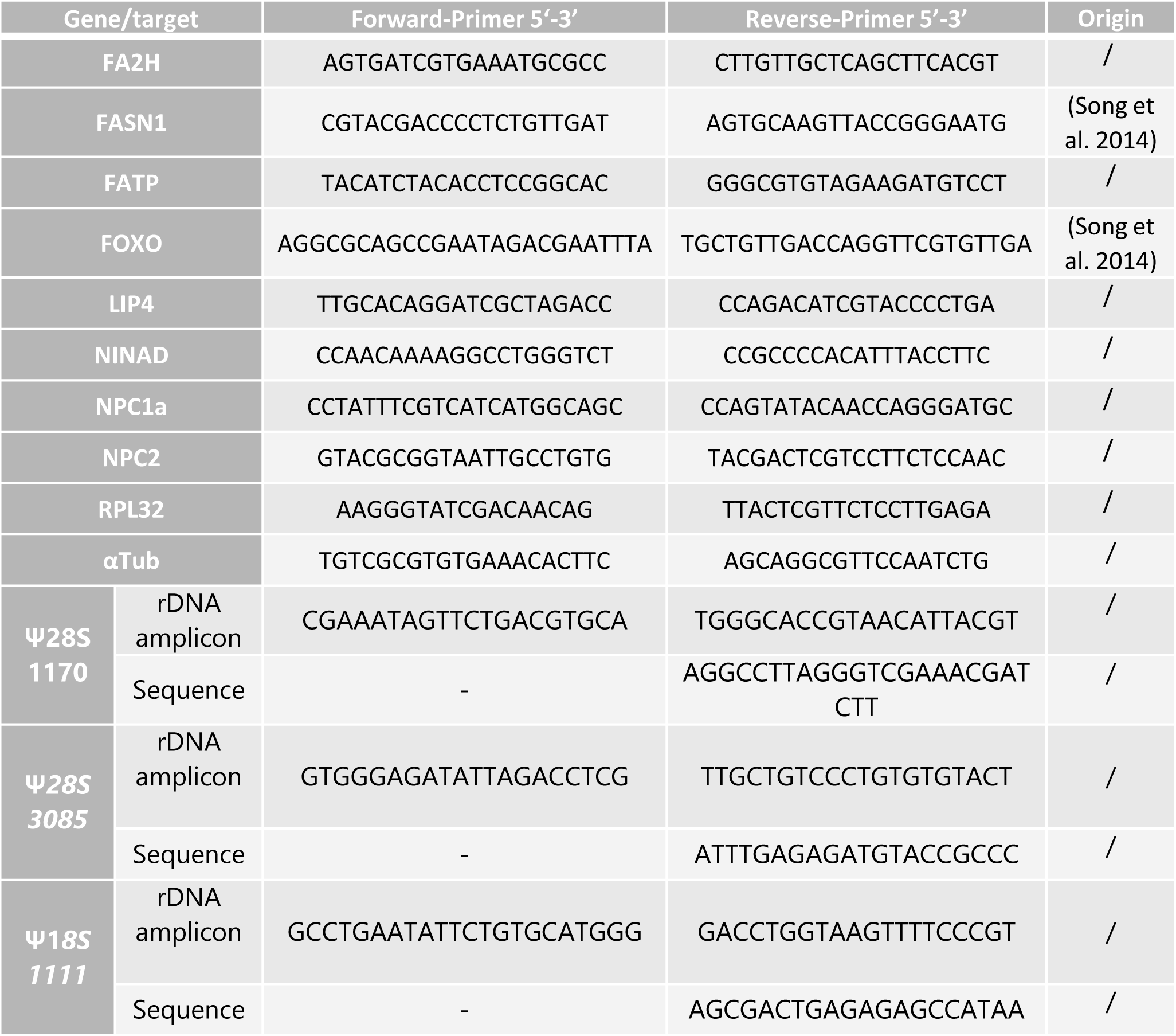
Sequences of primers used for qPCR and PSI-blot. The “gene/target” column represents the official names of the genes according to FlyBase or the position on the rDNA targeted and its purpose. The “Forward Primer 5’-3’ and “Reverse Primer 5’-3’ column show the primer sequences. The “Origin” column represents the source article of the primer, “/” signifies that the primer was conceived in the lab.

### Brain histology

For the paraffin section, flies were filled in collars and fixed for 4h in fresh Carnoy’s fixative (ethanol: chloroform: acetic acid at 30:15:5) as in Soulé & Martin (2020). Quantifications of neurodegenerative lesions were performed on the entire brains using ImageJ software. Data are expressed as number of vacuolar/lesions per µm^2^ of measured surface.

### Triglycerides measurement

The Triglycerides (TG) quantification has been performed with a colorimetric method using the enzymatic kit “LiquiColor TG kit” (Stanbio, Cat. #2200-430) as described in Soulé & Martin (2020). The TG determination was performed on 24 individuals of 7-day-old female fly.

### Sterols measurement

Sterols (including the cholesterol) in flies were measured as described in Soulé and Martin (2020). Measurements were carried out on 24 single 7-days-old adult female flies. Free sterol and sterol-ester levels were measured using the Amplex Red cholesterol assay kit (*Invitrogen*, Cat. #A12216), according to the manufacturer’s instructions. Notice that this enzymatic based-assay (kit) also measures all other sterols, and not solely the cholesterol (Serrano et al., 2024).

### CMCT-RT Pseudouridylation assay

Total RNAs from the different genotypes were extracted from the dissected gut with the SV Total RNA Isolation System Kit (Z3100, Promega), followed by a DNase-1 treatment (30 min at 37°C) to completely remove any DNA trace. Then the total RNA was purified with the NucleoSpin RNA Clean-up kit (740948.50, Macherey-Nagel), and the concentration determined. For pseudouridylation experiments, we follow the protocol of Huang et al. (2016), with some light modifications (Soulé et al., 2020). Dried total RNA (10µg for each reaction) were treated with (or without for control) N-cyclohexyl-N’-(2-morpholinoethyl)-carbodiimide metho-p-toluenesulphonate (CMC) for 30 min at 40°C, and subjected to alkali hydrolysis in the presence of 50 mM Na2CO3 (pH 10.4) at 50°C during 2 hours with shaking every 10 minutes. The primers of each probes (18S and 28S-rRNA) were radiolabeled using the T4 Polynucleotides Kinases (T4-PNK) and γ-P32-ATP. The 20 mers primers (see Suppl. Materials for the sequences) were designed about 50 nucleotides downstream to the predicted sites of pseudouridylation. The Reverse Transcription reactions were then performed on the 18S and 28S-rRNA templates respectively. In meantime, the DNA templates of the 18S and 28S-rRNA were generated by PCR using two primers located about 200 nucleotides upstream and downstream to the predicted pseudouridylation sites (a fragment of about 400 bp) (see Suppl. Materials for the sequences). For the sequencing of the DNA used as reference, we used the Sanger method and the Sequenase Kit (USB, Life Technologies), with slight modifications, notably because we used γ-P32 5’-labelled primers (the same as for the pseudouridylation reaction) instead of labeled nucleotides. Then, both reactions were migrated in parallel on an 8% acrylamide gel + 8M urea, at 2000 volts during 2 hours. After drying, the gels were exposed overnight, and then subjected to phosphoimager analysis using a Typhoon Trio Imaging System (GE Healthcare).

### HydraPsiSeq

The HydraPsiSeq experiments have been performed using the protocol as described in Marchand et al., (2020). In brief, total RNA (50-300 ng) from 7 day-old Drosophila’s dissected gut was subjected to hydrazine treatment (50% final concentration) for 45 min on ice. The reaction was stopped by ethanol precipitation using 0.3M NaOAc, pH5.2 and Glycoblue and incubated 30 min at -80°C. After centrifugation, the pellet was washed twice with 80% ethanol and re-suspended in 1M aniline, pH4.5. The reaction was incubated in dark for 15 min at 60°C and processed as described above, by ethanol precipitation.

RNA fragments obtained by hydrazine/aniline treatment were dephosphorylated at the 3’-end for 6h at 37°C using 10 U of T4 PNK in 100 mM Tris-HCl pH6.5, 100 mM MgOAc and 5 mM β-mercaptoethanol. T4 PNK was inactivated by incubation for 20 min at 65°C. RNA was extracted by phenol:chloroform and ethanol precipitated. The pellet was re-suspended in RNase free water. 3’-dephosphorylated RNA fragments were converted to library using the NEBNext® Small RNA Library Prep Set for Illumina® (NEB ref E7330S, USA) following the manufacturer’s recommendations. DNA library was quantified using a fluorometer (Qubit 3.0 fluorometer, Invitrogen, USA) and qualified using a High Sensitivity DNA chip on Agilent Bioanalyzer 2100. Libraries were multiplexed and subjected for high-throughput sequencing on an Illumina HiSeq1000 instrument with a 50 bp single-end read mode.

High quality raw sequencing reads (> Q30) were subjected to trimming using Trimmomatic v0.39 (Bolger et al., 2014) with the following parameters: MINLEN:08, STRINGENCY:7, AVGQUAL:30, trimmed reads were further aligned to the Drosophila rRNA (or rRNA/tRNA) reference sequence (accession number: PRJEB111190) using bowtie2 v2.4.4 (Langmead & Salzberg, 2012) in end-to-end mode (--no-unal --no-1mm-upfront -D 15 -R 2 -N 0 -L 10 -i S,1,1.15 as other bowtie2 parameters), only uniquely mapped reads in positive orientation were retained for further analysis. Mapped and sorted *.bam file was transformed to *.bed format. Locations of 5’-extremities of mapped unique reads were counted from *.bed file, giving raw cleavage profile. Reads’ 5’-end counts were normalized to local background in rolling window of 10 nucleotides, values for U residues were excluded to calculate the median. Locally normalized U profiles (NormUcount) were further transformed to U cleavage profiles only, by omitting values for other nucleotides. Resulting U profiles were used for calculation of “RiboMethSeq-like” scores (ScoreMEAN, A, B and PsiScore, equivalent to ScoreC in RiboMethSeq). Window of 4 neighboring nucleotides (+/-2 nt) was used for score calculation (Pichot et al., 2020). ScoreMEAN for each position is calculated in two steps, as follows: first, a ratio for number of cumulated 5’/3’-reads ends between preceding and following position is defined and, second, ScoreMEAN is calculated as a ratio of a drop for a given position compared to the average and variation for 4 neighboring positions (-2/+2). PsiScore (ScoreC2 in RiboMethSeq) is calculated using the formula :

PsiScore = 1-n_i_/(0.5*(SUM(n_j_*W_j_)/SUM(W_j_)+SUM(n_k_*W_k_)/SUM(W_k_)), where n_i_ – cumulated 5’-end U count for a given position, j – varies from i-2 to i-1, k varies from i+1 to i+2, Weight parameters are defined as 1.0 for -1 /+1 and 0.9 for -2/+2 positions. If not available, potential Psi positions in RNA sequence were predicted using a medium stringency combination of score mean2 >0.95 and ScoreA2>0.5, this combination generally gives the best balance between the sensitivity and selectivity in *de novo* detection. Quantification of Psi residues was done using PsiScore (similar to scoreC2 used in RiboMethSeq (Birkedal, et al., 2015; Pichot et al., 2020). This score is used for quantification of the modification level since it keeps linear calibration curve between protection and molar ratio of Psi residue in RNA.

### Puromycin experiments

Following dissection of 7 days old flies, guts were incubated 30 minutes at 25°C in Ringer’s buffer with 1mM of puromycine (ant-pr-1, invivoGen). Guts were then fixated in 4% PFA for two hours at 4°C. Then after washing with PBS + Triton 0,1%, they were incubated in anti-puromycine antibody (1:200) (mouse monoclonal 3RH11, Kerafast/Vectors Lab Company) overnight. After washing, the gut were incubated in anti-mouse alexa-488 antibody during 2 hours at 4°C. Gut were then mounted in Mowiol 4-88 (Calibochem, 475904) and images were recorded with Leica DM 600B microscope. Images of mid-gut were taken with the digital camera Hamamatsu C10600 ORCA-R (Leica). The fluorescence intensity was measured on ten enterocytes for each image with two images per gut (thus: n=20 enterocytes/gut) and between 20 to 30 guts/genotype, using ImageJ software.

### Quantification of the amount of ribosomes

Myo-Gal4>UAS-RPL10-GFP was used to express RPL10 ribosomal protein fused with GFP (Huang et al., 2013) in enterocytes. Following dissection of 7 days old flies’ guts were fixated in 4% PFA for two hours at 4°C. Then after washing with PBS + Triton 0,1%, gut were then mounted in Mowiol 4-88 (Calibochem, 475904) and images were recorded with Leica DM 600B microscope. Images of mid-gut were taken with the digital camera Hamamatsu C10600 ORCA-R (Leica). The fluorescence intensity was measured on ten enterocytes for each image with two images per gut (thus: n=20 enterocytes/gut) and between 20 to 30 guts/genotype, using ImageJ software.

### TRAP assay

Myo-Gal4>UAS-RPL13a-FLAG was used to express RPL13a ribosomal protein fused with FLAG (Huang et al., 2019) in enterocytes. Guts of 7 days old flies were dissected and homogenised in RIP Buffer [HEPES 20 mM, pH 7,4, KCl 300mM, Igepal® 0,75% (I8896, Sigma Aldrich), DTT 1mM, Cycloheximide 100µg/mL, MgCl2 5mM, Inhibitor of RNase SUPERase-IN™ (AM2694, Invitrogen) 250 U/mL and cOmplete™ Mini EDTA free (#11836170001, Roche)] inspired by (Huang et al., 2019). Homogenized guts were incubated 20 minutes at 4°C before centrifugation to remove cellular debris. 1/3 of the supernatant was stored at 80°C, the remaining supernatant was incubated with affinity gel ANTI-FLAG® M2 EZview™ (F2426, Millipore) for 90 minutes at 4°C. The affinity gel was then centrifugated and cleaned with RIP buffer 5 times. After cleaning, TRIzol (Invitrogen) was added to the gel to denature the ribosomes and release the translating RNA. After centrifugation, supernatant was kept and use to extract and purify the mRNA using standard TRIzol extraction followed by DNase treatment (Thermofisher, ref. EN0521) and a phenol/chloroform purification. Similar RNA extraction was performed on the 1/3 of supernatant that was previously isolated as a total RNA control. RNAs were then submitted to RT-qPCR using SybrGreen reagent (A25742, Applied Biosysthems™, ThermoFisher). qPCR results were normalized by the total RNA in order to investigate only the impact of the ribosome accessibility for the mRNA and eliminate the mRNA expression part.

### Statistical analyses

Statistical comparisons were done with GraphPad Prism. Data were analyzed using the student-T test. For the TRAP, two-fold ANOVA was used. Significance levels in figures were represented as *p < 0.05, **p < 0.01, ***p < 0.001. All quantitative data are reported as the mean ± S.E.M. (Standard Error of the Mean). Lifespan assays were subjected to survival analysis (log-rank test) using the freely available OASIS software (Yang et al., 2011). See Table 2 for the detailed statistics.

**Table 2.** Longevity Statistics. The « Genotype » column represents the genotype of the flies. The “Number” column represents to total number of flies tested. The “Age at % mortality” column represents the day at which 50% and 100% of the flies were dead. The “Statistics” column represents with which genotype the flies were compared, and the log-rank test p-values calculated by OASIS 2 Software (Yang et al., 2011).

| Genotype | Number | Age at % mortality |  | Statistics |  |
| --- | --- | --- | --- | --- | --- |
|  |  | 50% | 100% | Compared with genotype | p-Value |
| WT | 192 | 28 | 32 | - | - |
| Jou- | 244 | 32 | 42 | WT | P<0,0001 |
| sno2- | 259 | 30 | 44 | WT | P<0,0001 |
| sno3- | 258 | 32 | 42 | WT | P<0,0001 |
| sno2&3- | 272 | 18 | 37 | WT | P<0,0001 |
| jou-, Mex-GS > jou-, UAS-jou -RU | 256 | 27 | 69 | - | - |
| jou-, Mex-GS > jou-, UAS-jou +RU | 231 | 23 | 35 | jou-, Mex-GS > jou-, UAS-jou -RU | P<0,0001 |
| sno2-, Mex-GS > sno2-, UAS-sno2 -RU | 134 | 64 | 74 | - | - |
| sno2-, Mex-GS > sno2-, UAS-sno2 +RU | 161 | 45 | 61 | sno2-, Mex-GS > sno2-, UAS-sno2 -RU | P<0,0001 |
| Sno3-, Mex-GS > sno3-, UAS-sno3 -RU | 221 | 58 | 76 | - | - |
| Sno3-, Mex-GS > sno3-, UAS-sno3 +RU | 195 | 43 | 71 | Sno3-, Mex-GS > sno3-, UAS-sno3 -RU | P<0,0001 |
| sno2&3-, Mex-GS > sno2&3-, UAS-sno2&3 -RU | 288 | 36 | 66 | - | - |
| sno2&3-, Mex-GS > sno2&3-, UAS-sno2&3 +RU | 140 | 44 | 80 | sno2&3-, Mex-GS > sno2&3-, UAS-sno2&3 -RU | P<0,0001 |

## Acknowledgments

We thank L. Mellottée for her technical assistance in the genesis of the UAS-snoRNA. H. Tricoire (Paris, France), and B.A. Edgar (Heidelberg, Germany) for fly stocks. This work was supported by the ANR (Agence Nationale de la Recherche, Ageing-jou, France), and by the CNRS (France).

## Author Contributions

JRM conceived the experiments. JRM and TG designed the experiments. TG and PM perform the experiments and analyzed data. VM and YM performed the HydraPsiSeq. JRM wrote the manuscript with input from all authors.

## Supplementary Information

Two Supplementary Tables are available online.

## Data Availability

Raw HydraPsiSeq data are available at the European Nucleotide Archive (https://www.ebi.ac.uk/ena/browser/home) under the accession number PRJEB111190. The authors declare that all other data and the methods used in this study are available within this article. Supplementary Information is available from the corresponding authors upon reasonable request.

## References

Abe, M & Bonini, NM (2013). MicroRNAs and neurodegeneration: role and impact. Trends in Cell Biology, 23, 30–36.

Bailey AP et al. (2015). Antioxidant Role for Lipid Droplets in a Stem Cell Niche of Drosophila. Cell 163, 340–353.

Beller, M et al. (2010). PERILIPIN-Dependent Control of Lipid Droplet Structure and Fat Storage in Drosophila. Cell Metabolism, 12, 521–532.

Benes, H et al. (1996). Overlapping Lsp-2 gene sequences target expression to both the larval and adult Drosophila fat body. Insect Mol Biol, 5, 39–49.

Bi, J et al. (2012). Opposite and redundant roles of the two Drosophila perilipins in lipid mobilization. Journal of Cell Science, 125, 3568–3577.

Biglou, SG, Bendena, WG, & Chin-Sang, I (2021). An overview of the insulin signaling pathway in model organisms Drosophila melanogaster and Caenorhabditis elegans. Peptides, 145, 170640.

Birkedal, U, et al., (2015). Profiling of ribose methylations in RNA by high-throughput sequencing. Angew Chem Int Ed Engl, 54, 451–455.

Bolger, AM, Lohse, M, & Usadel, B (2014). Trimmomatic: a flexible trimmer for Illumina sequence data. Bioinformatics, 30, 2114–2120.

Bolukbasi E et al. (2017). Intestinal Fork Head Regulates Nutrient Absorption and Promotes Longevity. Cell Rep, 21, 641–653.

Broughton, S.J. et al. (2005). Longer lifespan, altered metabolism, and stress resistance in Drosophila from ablation of cells making insulin-like ligands. Proceedings of the National Academy of Sciences of the United States of America, 102, 3105–3110.

Carvalho M, et al. (2010). Survival strategies of a sterol auxotroph. Development, 137, 3675–3685.

Charrette, M and Gray, MW (2000). Pseudouridine in RNA: What, Where, How, and Why. IUBMB Life, 49: 341–351.

Diaz G et al. (2008). Hydrophobic characterization of intracellular lipids in situ by Nile Red red/yellow emission ratio. Micron (Oxford, England: 1993), 39, 819–824.

Dutta D et al. (2015). Regional Cell-Specific Transcriptome Mapping Reveals Regulatory Complexity in the Adult Drosophila Midgut. Cell Rep, 12, 346–358.

Eacker SM, Dawson TM, & Dawson VL (2009). Understanding microRNAs in neurodegeneration. Nature Rev Neurosci, 12, 837–741.

Fontana L, Partridge & Longo VD (2010). Extending healthy life span--from yeast to humans. Science, 328, 321–326.

Gems D, & Partridge L (2013). Genetics of longevity in model organisms: debates and paradigm shifts. Annual Review of Physiology, 75, 621–644.

Giannakou ME & Partridge L (2007). Role of insulin-like signalling in Drosophila lifespan. Trends in Biochemical Sciences, 32, 180–188.

Graveley BR et al. (2011). The developmental transcriptome of Drosophila melanogaster. Nature 471, 473–479.

Haussmann IU, et al., (2022). CMTr cap-adjacent 2’-O-ribose mRNA methyltransferases are required for reward learning and mRNA localization to synapses. Nat Commun, 13, 1209.

Heier C & Kühnlein, RP (2018). Triacylglycerol Metabolism in Drosophila melanogaster. Genetics, 210, 1163–1184.

Heiman, M, et al. (2014) Cell type–specific mRNA purification by translating ribosome affinity purification (TRAP). Nature Protocols, 9, 1282–1291.

Huang, et al. (2013) Translational Profiling of Clock Cells Reveals Circadianly Synchronized Protein Synthesis. PLoS Biology, 11(11): e1001703.

Huang, K et al. (2019) RiboTag translatomic profiling of Drosophila oenocytes under aging and induced oxidative stress. BMC Genomics, 20, 50.

Huang X, Warren JT, Buchanan J, Gilbert LI, & Scott MP (2007). Drosophila Niemann-Pick Type C-2 genes control sterol homeostasis and steroid biosynthesis: a model of human neurodegenerative disease. Development, 134, 3733–3742.

Jack K, et al., (2011). rRNA Pseudouridylation Defects Affect Ribosomal Ligand Binding and Translational Fidelity From Yeast to Human Cells. Mol Cell, 44, 660–666.

Jaque-Cabrera L, Buggiani J, Bignon J, Daira P, Bernoud-Hubac N, & Martin JR (2025). Knockdown of the snoRNA-Jouvence Blocks the Proliferation and Leads to the Death of Human Primary Glioblastoma Cells. Noncoding RNA, 11(4):54.

Jiang H et al. (2009). Cytokine/Jak/Stat signaling mediates regeneration and homeostasis in the Drosophila midgut. Cell, 137, 1343–1355.

Jing X & Behmer ST (2020). Insect Sterol Nutrition: Physiological Mechanisms, Ecology, and Applications. Annual Review of Entomology, 65, 251–271.

Johnson AA, & Stolzing A (2019). The role of lipid metabolism in ageing, lifespan regulation, and age-related disease. Ageing Cell, 18, e13048 (2019).

Kato M & Slack FJ (2013). Ageing and the small non-coding RNA world. Ageing Res Rev., 12, 429–435.

Kiss T (2002). Small nucleolar RNAs: an abundant group of noncoding RNAs with diverse cellular functions. Cell, 109, 145–148.

Langmead, B, & Salzberg, SL (2012). Fast gapped-read alignment with Bowtie 2. Nat Methods 9, 357–359.

Liu L et al. (2015). Glial lipid droplets and ROS induced by mitochondrial defects promote neurodegeneration. Cell, 160, 177–190.

Liu N et al. (2012). The microRNA miR-34 modulates ageing and neurodegeneration in Drosophila. Nature, 482, 519–523.

López-Otín C, Blasco MA, Partridge L, Serrano M, & Kroemer G (2013). The hallmarks of ageing. Cell, 153, 1194–217.

López-Otín, C et al. (2016). Metabolic Control of Longevity. Cell, 166, 802–821.

López-Otín, C et al. (2023). Hallmarks of ageing: An expanding universe. Cell, 186, 243–278.

Marchand, V et al., (2020). HydraPsiSeq: A Method for Systematic and Quantitative Mapping of Pseudouridines in RNA. Nucleic Acids Research, 48 (19): e110-e110.

Morgan NS, Skovronsky DM, Artavanis-Tsakonas S & Mooseker MS (1994). The molecular cloning and characterization of Drosophila melanogaster myosin-IA and myosin-IB. J. Mol. Biol., 239, 347–356.

Ofengand, J and Bakin, A (1997) Mapping to nucleotide resolution of pseudouridine residues in large subunit ribosomal RNAs from representative eukaryotes, prokaryotes, archaebacteria, mitochondria and chloroplasts. Journal of Molecular Biology, 266, 2, 246–268.

Owusu-Ansah E, & Perrimon N (2014). Modeling metabolic homeostasis and nutrient sensing in Drosophila: implications for ageing and metabolic diseases. Disease Models & Mechanisms, 7, 343–350.

Phillips SE, Woodruff EA, Liang P, Patten PM, & Broadie K (2008). Neuronal loss of Drosophila NPC1a causes cholesterol aggregation and age-progressive neurodegeneration. J Neurosci., 28, 6569–6582.

Pichot, F, Marchand, V, Ayadi, L, Bourguignon-Igel, V, Helm, M, & Motorin, Y (2020). Holistic Optimization of Bioinformatic Analysis Pipeline for Detection and Quantification of 2’-O-Methylations in RNA by RiboMethSeq. Front Genet, 11, 38.

Poirier L, Shane A, Zheng J, & Seroude L (2008). Characterization of the Drosophila Gene-Switch system in ageing studies: a cautionary tale. Ageing Cell, 7, 758–770.

Saher G (2023). Cholesterol Metabolism in Ageing and Age-Related Disorders. Annual Review of Neuroscience, 46, 59–78.

Schmidt, EK et al. (2009). SUnSET, a nonradioactive method to monitor protein synthesis. Nat Methods, 6:275–277. doi: 10.1038/nmeth.1314.

Serrano J et al. (2024). The importance of choosing the appropriate cholesterol quantification method: enzymatic assay versus gas chromatography. Journal of Lipid Research, 65, 100561.

Singh PP, Demmitt BA, Nath RD, & Brunet A. (2019). The Genetics of Ageing: A Vertebrate Perspective. Cell, 177, 200–220.

Soulé S, Mellottée L, Arab A, Chen C, & Martin JR (2020). Jouvence a small nucleolar RNA required in the gut extends lifespan in Drosophila. Nat. Commun., 11, 1–21.

Soulé S, & Martin JR (2020). ninaD regulates cholesterol homeostasis from the midgut which protects against neurodegeneration. BioRxiv (doi.org/10.1101/2020.11.27.401059).

Tschäpe JA, Hammerschmied C, Mühlig-Versen M, Athenstaedt K, Daum G, & Kretzschmar D (2002). The neurodegeneration mutant lochrig interferes with cholesterol homeostasis and Appl processing. EMBO J., 21, 6367–6376.

Voght SP, Fluegel ML, Andrews LA, & Pallanck LJ (2007). Drosophila NPC1b Promotes an Early Step in Sterol Absorption from the Midgut Epithelium. Cell Metab., 5, 195–205.

Yang JS et al. (2011). OASIS: Online Application for the Survival Analysis of Lifespan Assays Performed in Ageing Research. PLoS ONE, 6, e23525.

Ye K (2007). H/ACA guide RNAs, proteins and complexes. Curr Opin Struct Biol., 17, 287–292.

You, S et al. (2021) Circadian regulation of the Drosophila astrocyte transcriptome. PLoS Genetics, 17(9): e1009790.

